# Is the whole more than the sum of its parts? Considering global and local features of the connectome improves prediction of individuals and phenotypes

**DOI:** 10.1101/2025.10.22.683965

**Authors:** Steve Riley, Annie Cheng, Yu-Wei Wang, Xilin Shen, Yize Zhao, Avram Holmes, R. Todd Constable, Sarah W. Yip

**Affiliations:** Department of Psychiatry, Yale School of Medicine, New Haven, CT, USA, United States; Radiology and Biomedical Imaging, Yale University School of Medicine, New Haven, CT, USA, United States; Department of Biostatistics, Yale School of Public Health, Yale University, New Haven, CT, USA, United States; Department of Psychiatry, Brain Health Institute, Rutgers University, Piscataway, NJ, United States; Child Study Center, Yale School of Medicine, New Haven, CT, USA, United States

## Abstract

Popular methods for analyzing the brain’s functional connectome examine statistical associations between pairs of atlas-defined brain regions, viewing the strength of these links as independent values. However, edges within a standard connectivity matrix, i.e., correlations between individual regions or nodes, are not independent. They are part of an interconnected system. Here, we propose that consideration of both independent, linear relationships (as in standard approaches such as linear kernel ridge regression and connectome-based predictive modeling) as well as higher order statistical associations - such as tertiary interactions between matrix components and global features of the matrix space - will enhance identification of meaningful individual differences. To test this, we adopt a geometrically grounded measure of similarity that accounts for higher-order local statistical relationships and global interactions, the Wasserstein metric. Results indicate that considering connectivity matrices as representations of their associated Gaussian distributions significantly improves both identification of individuals based on their connectivity matrices (aka, ‘fingerprinting’) and prediction of individual differences in phenotypes such as fluid intelligence and openness to experience. Thus, both pairwise local and global brain connectivity properties encode for meaningful individual differences that relate to phenotypic expressions and should be considered in brain-behavior predictive models.

## Introduction

Identifying reliable associations between individual differences in the brain and individual differences in personality, behaviors and clinical presentations is a fundamental goal of basic and clinical neuroscience. In recent years, work in this area has leveraged advances in machine learning to build brain-based predictive models of cognitive abilities such as IQ and attention - and of psychiatric variables such as illness severity and treatment outcomes - using functional connectivity matrices (aka ‘connectomes) [1-8]. However, despite numerous strengths, higher precision, accuracy and increased explained variance in predicting outcomes are still needed for both real world translational and clinical applications, and to improve understanding of basic brain mechanisms. While multiple methods for predictive brain-behavior modeling exist, in general these approaches focus on pairwise relationships (i.e., edges in a vectorized connectivity matrix) and variation between these approaches tends to be at the algorithmic level, for example using different types of penalized regression to boost feature selection [3, 9-11]. Here, we test an alternative approach by asking whether accounting for additional information in the matrix space might improve model performance and ask what the implications of that might mean for our understanding of the human brain.

Common methods for analyzing functional connectomes typically examine statistical or informational associations between pairs of atlas-defined brain regions, viewing the strength of these links as independent values [12-17]. However, edges within a standard connectivity matrix, i.e., correlations between individual regions or nodes, are not independent [18] - they are part of an interconnected system - in other words, each pairwise value imposes constraints on all other pairwise values. However, this inter-relatedness (the existence of higher order statistical associations) is not accounted for by common contemporary brain-behavior modeling approaches such as standard connectome-based predictive modeling or kernel ridge regression approaches. Here, we propose that considering connectivity matrices as representations of their associated Gaussian distribution enables consideration of both independent, linear relationships (as in standard approaches) as well as higher order statistical associations, such as tertiary interactions between matrix components and global features of the matrix space. In doing so, we further hypothesized that this consideration of both local- *and* systems-level statistical relationships would outperform models built using either information source alone.

To test these theories, we selected a global, geometrically grounded measure of similarity that accounts for higher-order local statistical relationships and global interactions among brain regions - the Wasserstein distance metric. As shown in Figure 1, unlike angular distance (which is akin to correlation, details in Supplement) [14] and other non-Wasserstein metrics tested here, the *p* -Wasserstein metric quantifies distance over the space of probability distributions. In doing so, it circumvents the need to first vectorize the matrix (as in standard common approaches), and therefore retains higher order statistical interactions between edges that are lost in the vectorization process, enabling feature selection within the full matrix space. As further illustrated in Figure 1, the *p* -Wasserstein distance (as rooted in optimal transport theory), defines the distance between two distributions as the minimal cost of transporting mass to transform one distribution into another, where the cost of moving one unit of mass equals the distance it is moved with respect to the *p* -norm. Conceptually, this can be thought of as the amount of work needed to transform a given pile of dirt into a different pile of dirt (**Figure 1**) [19]. While not previously used in brain-behavior modeling, the Wasserstein metric is employed in other Bayesian approaches and in other machine learning applications due to its ability to capture subtle differences between distributions [20, 21].

**Figure 1.**
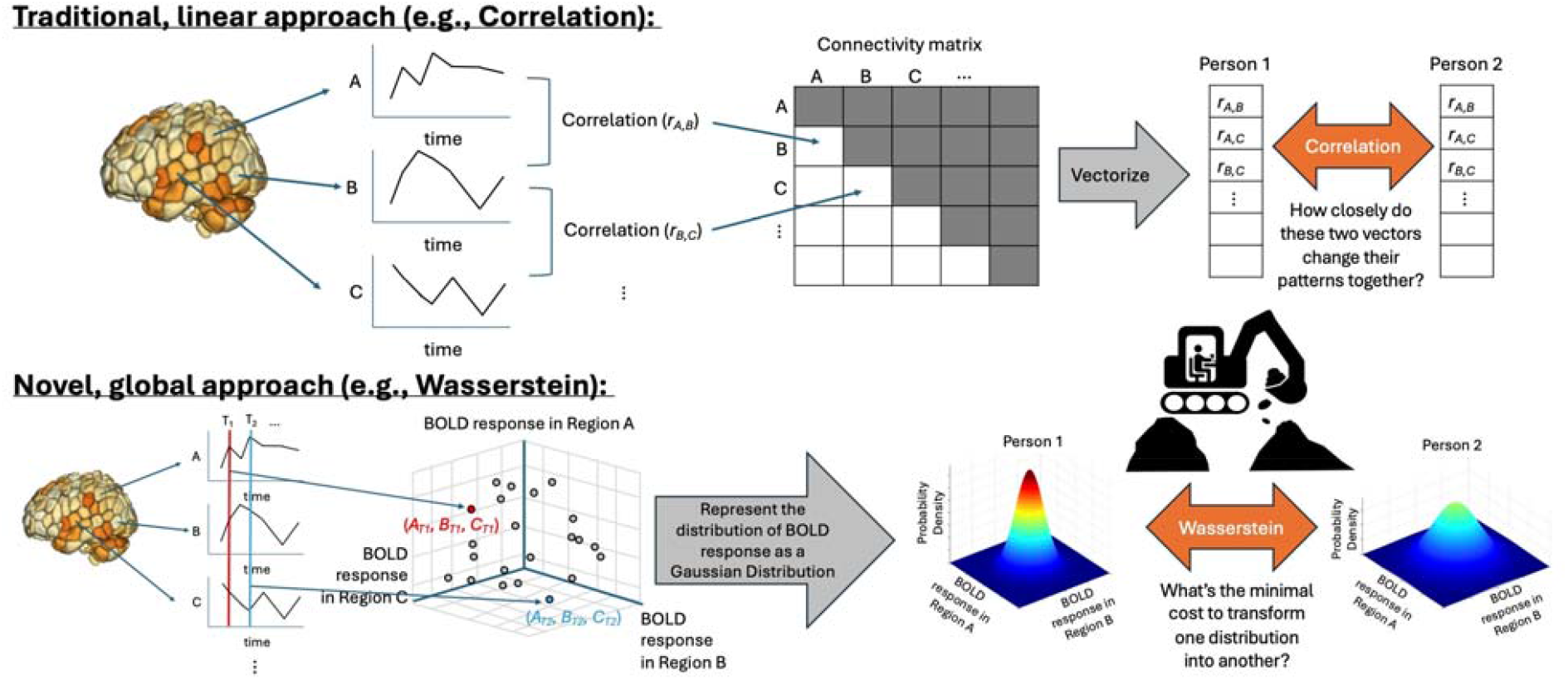
Schematic diagram detailing example of a traditional, linear approach versus the proposed approach. A linear correlation approach involves computing pairwise correlations among all brain regions and vectorizing the connectivity matrix, which thereby treats the edges (i.e., strength of association between brain regions) as independent values; by contrast, the global, Wasserstein approach represents each functional connectivity matrix as a shared, Gaussian distribution, which enables consideration of both independent, linear relationships and also higher-order statistical associations among the edges. Conceptually, the Wasserstein may be considered as summarizing the amount of work needed to transform a given pile of dirt into a different pile of dirt, and is therefore sometimes referred to as the ‘earth mover’s distance metric [19].

We first tested the ability of the Wasserstein metric to correctly identify or match separate connectivity matrices obtained from the same individual during performance of different cognitive tasks, commonly referred to as ‘fingerprinting’ or ‘ID rate’[1]. We hypothesized that ID rate conducted using the Wasserstein metric would have higher overall accuracy (more successful ID’s) versus ID rate conducted using standard correlation, thereby indicating that considering matrices as representations of their associated Gaussian distributions would enable retention of connectivity features that meaningfully differ between individuals, and which are not captured when using standard vector-based approaches. We next tested the utility of the Wasserstein metric to improve brain-behavior predictive models generated using functional connectomes. To do so, we developed a pipeline that allows for flexible incorporation of different metrics and compared the Wasserstein’s utility against other standard distance metrics, such as angular and Euclidean distance, and versus two other brain-behavior modeling approaches: connectome-based predictive modeling (CPM) and kernel ridge regression (KRR). We again hypothesized that this consideration of functional connectivity matrices as representations of their associated Gaussian distributions would outperform standard, non-distribution-based approaches, as indicated by increased brain-behavior predictive accuracy.

In all cases, the Wasserstein approach outperformed other, non-distribution-based metrics and methods. Unlike other approaches, use of the Wasserstein metric to compare connectivity matrices simultaneously enables consideration of local pairwise interactions (i.e., connectivity between two nodes, as in existing standard approaches), of more complex local statistical interactions (e.g., three-way interactions between trios of nodes), and of global matrix properties (e.g., covariance in connectivity shared across all nodes). As a final test we therefore sought to determine which of these three features might be driving improved performance. Results indicated that both local pairwise and global properties contribute to model accuracy, demonstrating that both local and global brain connectivity properties encode for meaningful individual differences that relate to phenotypic expressions.

## Methods

### The Wasserstein metric

The *p* -Wasserstein metric, or Earth Mover’s Distance, provides a distance measure on the space of probability distributions. Rooted in optimal transport theory, the *p* -Wasserstein distance between two distributions is defined as the minimal cost of transporting mass to transform one distribution into another, where the cost of moving one unit of mass equals the distance it is moved with respect to the *p* -norm. For two probability measures *μ* and *v* on a metric space *X*, the *p* -Wasserstein distance *W*_*p*_ (μ, ν) is given by:

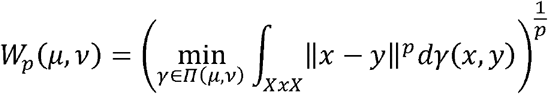

where *Π* (μ, ν) denotes the set of all joint distributions on *X* x *X* that have marginal distributions *μ* and *ν* [22].

In general, calculation of this distance is an intractable problem. However, the 2-Wasserstein distance between multivariate normal distributions admits a closed form [23]:

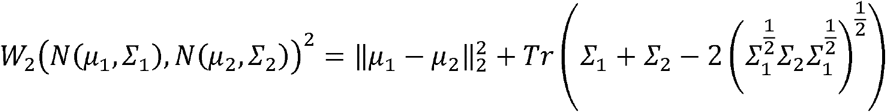

where *N* (*μ, Σ*) is the normal distribution with mean *μ* and covariance matrix *Σ, Tr(X*) is the trace of matrix *X*, and 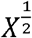 is the principal matrix square root of *X*. We associate an FC matrix *A* with *N* (0,*A*), which is the zero-mean normal distribution that has covariance matrix *A*. So, for our purposes, the Wasserstein distance between FC matrices *A* and *B* (also known as the Bures-Wasserstein metric) is:

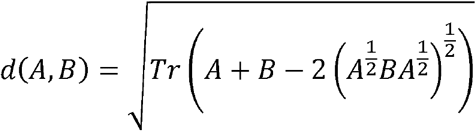

### Using the Wasserstein to ‘ID’ connectivity matrices across individuals and brain states

To test the initial hypotheses that consideration of both local and global properties of the matrix space, as encompassed via the Wasserstein metric, might improve standard connectivity-based approaches, we first tested the ability of this approach to correctly identify or match separate connectivity matrices obtained from the same individual at different times or during performance of different cognitive tasks. This well-established approach, commonly referred to as ‘fingerprinting’ or ‘ID rate’, follows the method from Finn *et al*. (2015), which showed that different functional connectivity (FC) matrices computed from fMRI data acquired from the same individual over different runs or acquisitions are similar enough that matrices from the same individual can be matched even among hundreds of “distractor” matrices from other participants [1]. This method of establishing the ID rate uses correlations between vectorized FC matrices as a similarity measure: if a participant’s matrix from scan x is more highly correlated with their matrix from scan y than it is with any other participant’s scan y, then the individual is considered to have been successfully identified, with ID rate quantified as the percentage of individuals in a given dataset who are successfully identified.

Using data from the Human Connectome Project (HCP), collected from 515 individuals over 9 different acquisitions (corresponding to data collected during two different resting state runs and seven different tasks/brain-states), we used functional connectivity matrices computed using the Shen 268-node atlas (details in Supplement). We then compared ID rate as quantified using angular distance (which is mathematically identical to using Pearson’s r, as in standard ID rate [1], see Supplement) against ID rate quantified using Wasserstein, Euclidean, and Manhattan distance metrics.

### Using the Wasserstein to improve accuracy of brain-behavior models

To test the utility of the Wasserstein metric in improving accuracy of brain-behavior models, we created a modified, flexible regression pipeline loosely based on CPM [1] with built-in cross-validation, referred to as connectome regression in any metric (CRAM, Figure 3). This approach uses multidimensional scaling (MDS, see Supplement for details on MDS) to reduce the dimension of the space of FC matrices and then applies linear regression to establish a predictive model of an outcome as a function of an FC matrix. Permutation testing in training data is then used to recover relevant features of FC matrices that are predictive of outcomes.

### Predictive modeling with CRAM

For a model to be predictive, it must be testable on data that is held out from the training step. While classic MDS has no method to incorporate new data points into the output space, an extension known as landmark MDS [24] does have this capability. Here, the initial transformation is computed based on a training set of data points that become ‘landmarks’ linking the original space to the new one. New data is then positioned by minimizing stress of their subsequent transformation. Thus, landmark MDS allows for the MDS and regression steps of the pipeline to be jointly cross validated, preventing data ‘leakage’ at either step.

The choice of dimension for the MDS step might affect the performance of CRAM. To circumvent the need to tune a hyperparameter at this step, we use a common ‘rule-of-thumb’ to keep all dimensions where the variance accounted for by that extra embedding dimension is greater than average. In PCA, a closely related technique to MDS, this choice would be equivalent to retaining all dimensions where the eigenvalue is greater than one [25]. Other methods exist for optimizing the dimension, such as cross-validation; however, any such method for tuning k must then be further validated on out-of-sample data, requiring a nested approach. We do not examine these other methods for choosing k to maintain simplicity and reduce computation time. As reported in Table S7 and Figure S3 in the Supplement, analyses using different dimensionality thresholds (ranging from 50% to 150% of the rule-of-thumb cutoff) provided comparable results, suggesting that model performance and feature identification are not critically dependent on the number of dimensions selected during MDS.

Once the predictive model is established, a question of interest is how to determine which connections in the FC matrix are driving the model predictions so that we may associate the model with relevant anatomical features. To answer this question, we employ permutation testing and *post hoc* feature selection. We create 1000 permuted models, indexed by *m*. A single permuted model at index *m* involves two steps: (1) creation of a randomly permuted set *y*_*m*_ of the actual outcome values *y* and (2) generation of a CRAM model ŷ_*m*_ *= f*_*m*_ (*X*), based on the permutation *y*_*m*_, that can predict an outcome measure ŷ_*m*_ for an arbitrary input matrix *X*. For each permuted model, we calculate the correlations (Pearson’s *r*) between entries at location *i,j* of the input matrices with their predicted outcomes ŷ_*m*_ . This generates a value *r* _*i,j,m*_ = *corr* (*A*_*i j*_, *f* _*m*_(*A*)) for each combination of *i, j*, and *m*. When aggregating across all *m* we now have a non-parametric null distribution of *r*_*i j*_ values for each location *i, j*.

Let ŷ = *f* (*A*) be the CRAM model for the original, unpermuted outcome data *y* . Let *R*_*i,j*_ = *corr* (*A*_*ij*_, *f* (*A*)). We determine that an edge at location *i, j* is associated with this model if it is outside the two-sided 95% interval of values obtained from the permuted CRAM models. So, if *R*_*i,j*_ ≥ *r*_*i j,m*_ for 97.5% of *m* values, then it is determined to be positively associated with the outputs of the true model. Similarly, this edge is negatively associated with the model if *R*_*i,j*_ ≤ *r*_*i j,m*_ for 97.5% of *m* values. Let *M* ^+^ be the matrix of positive edges: 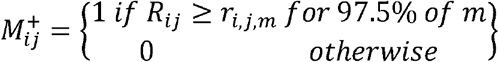 . Similarly, define the matrix *M* ^-^ of negative edges: 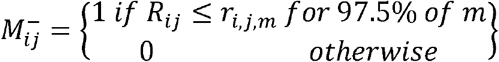 . Here, positive (negative) edges correspond to edges for which increased (decreased) connectivity is a positive predictor of the outcome of interest, as in standard CPM [26]. We test how different significance thresholds affect mask properties in the supplement.

### Feature stability

To prevent over-fitting to a random split of the data, current predictive modeling guidelines suggest iterating k-fold CV and averaging performance across iterations [27]. Here we use this approach to compare the stability of features selected across different iterations of CRAM using different metrics and compare this to the stability of features selected across iterations of CPM.

## Results

### Using the Wasserstein to ‘ID’ connectivity matrices across individuals and brain states

To test the initial hypotheses that consideration of both local and global properties of the matrix space, as encompassed via the Wasserstein metric, might improve standard connectivity-based approaches, we first tested the ability of this approach to correctly identify or match separate connectivity matrices obtained from the same individual at different times or during performance of different cognitive tasks. This well-established approach, commonly referred to as ‘fingerprinting’ or ‘ID rate’, follows the method from Finn *et al*. (2015), which showed that different functional connectivity (FC) matrices computed from fMRI data acquired from the same individual over different runs or acquisitions are similar enough that matrices from the same individual can be matched even among hundreds of “distractor” matrices from other participants [1]. This method of establishing the ID rate uses correlations between vectorized FC matrices as a similarity measure: if a participant’s matrix from scan x is more highly correlated with their matrix from scan y than it is with any other participant’s scan y, then the individual is considered to have been successfully identified, with ID rate quantified as the percentage of individuals in a given dataset who are successfully identified.

Using data from the Human Connectome Project (HCP), collected from 515 individuals over 9 different acquisitions (corresponding to data collected during two different resting state runs and seven different tasks/brain-states), we used functional connectivity matrices computed using the Shen 268-node atlas (details in Supplement). We then compared ID rate as quantified using angular distance —which is mathematically identical to using Pearson’s r, as in standard ID rate [1]—against ID rate quantified using Wasserstein, Euclidean, and Manhattan distance metrics. As shown in **Figure 2**, ID rate using the Wasserstein metric outperformed all other metrics across all rest and task acquisitions, with an overall Wasserstein ID rate of 93.6% vs 68.6% for angular distance (Figure S1). These results suggest that considering matrices as representations of their associated Gaussian distributions enables retention of connectivity features that meaningfully differ between individuals, and which are not captured when using standard vector-based approaches.

**Figure 2.**
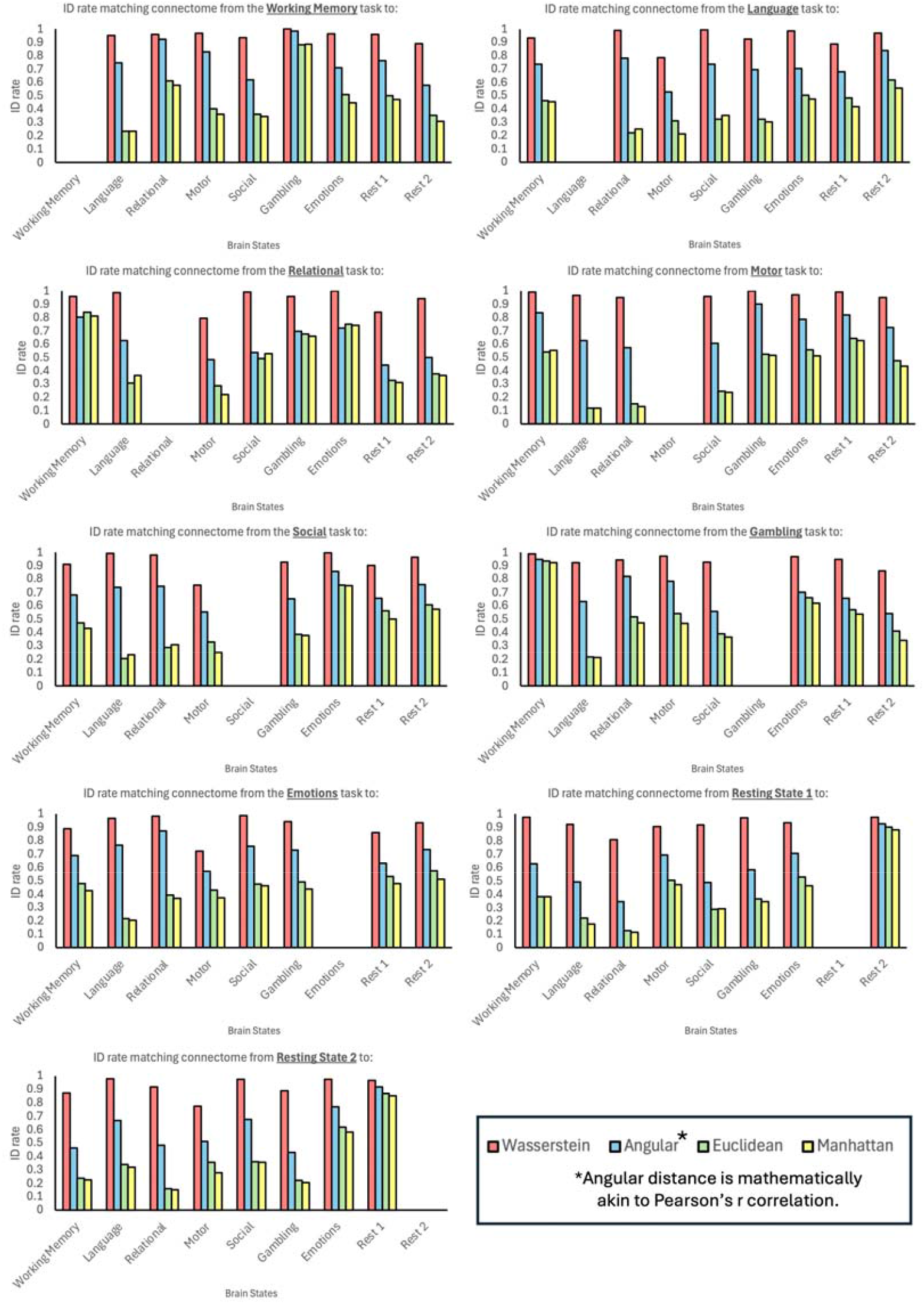
**Average ‘ID rate’** using the four metrics of interests across all possible pair-wise combination of connectivity matrices computed from the seven tasks and two resting state runs from the Human Connectome Project (HCP, N=515). ID rate is calculated per [1]. Findings show that a distribution-based metric, the Wasserstein distance metric, outperforms other distance metrics.

### Prediction of fluid intelligence using the Wasserstein

Having established that the Wasserstein metric outperforms other vector-based similarity measures in retaining meaningful information about individuals across scans, we next sought to test its sensitivity to improve performance of brain-behavior predictive models. In other words, we next asked whether the information retained by our Gaussian approach was also sufficiently meaningful to improve prediction of non-fMRI-derived phenotypes. We again used the two resting state and seven tasks from the HCP data set and tested how well we could predict fluid intelligence from FC matrices as measured by the Raven’s Matrices task, as in prior formative work [1].

To perform this test, we developed a pipeline that allows for flexible incorporation of different metrics using a combination of multidimensional scaling and ordinary least-squares linear regression, hereafter referred to as ‘connectome regression with any metric’ (CRAM, **Figure 3**, see Methods for details on multidimensional scaling and other model parameters). This allowed us to test brain-behavior model performance across the different distance metrics while keeping all other methods the same. To further benchmark performance against relatively widely used brain-behavior modeling approaches, we also conducted ‘standard’ connectome-based predictive modeling (CPM; i.e., vector-based linear feature selection using correlation, no multi-dimensional scaling) and kernel ridge regression (KRR). Models were run with ten-fold cross-validation and averaged over 100 independent runs to prevent over-fitting to a random ten-fold split [27], with permutation testing.

**Figure 3.**
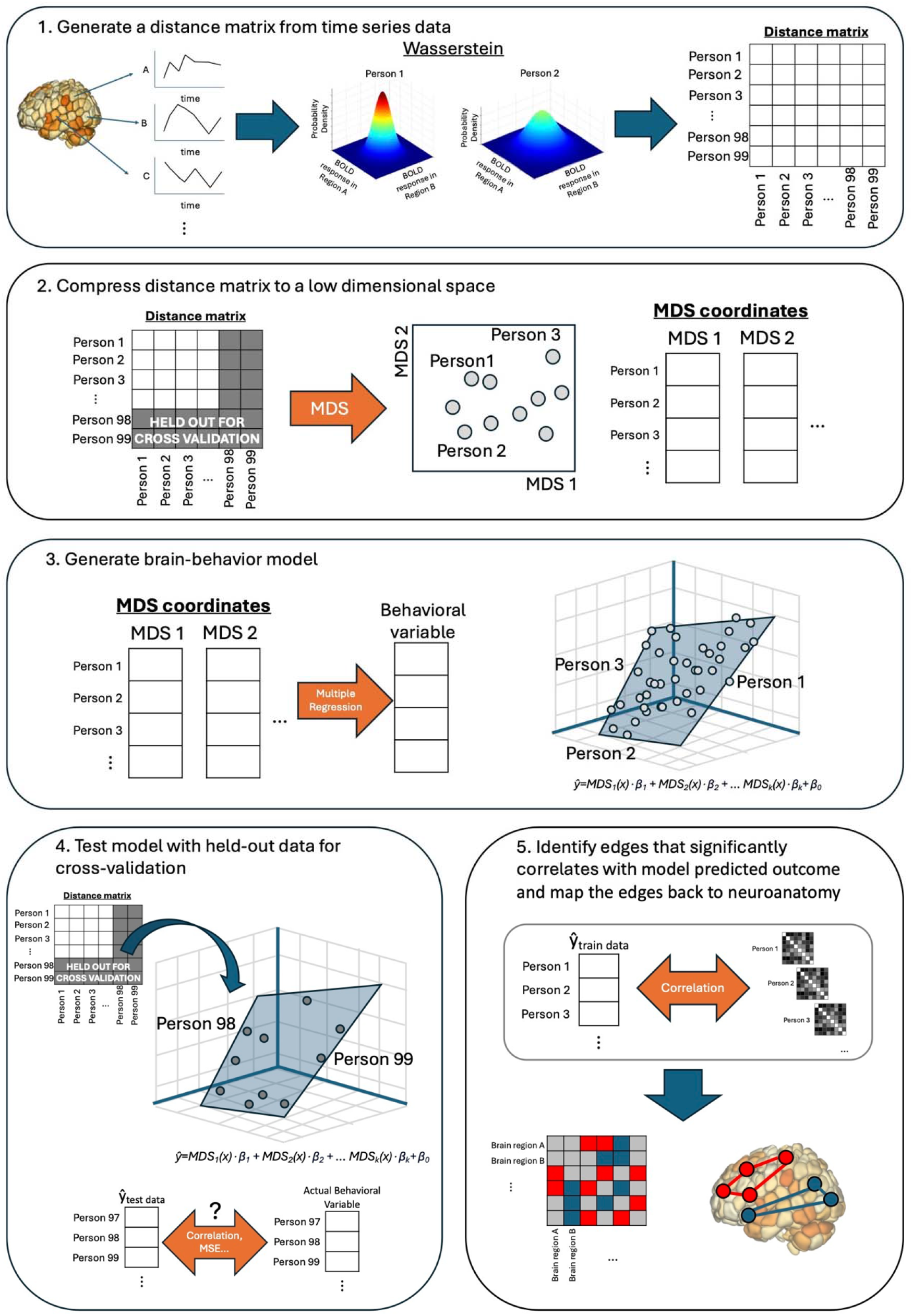
A schematic overview of connectome-regression with any metric (CRAM). During CRAM, (1) a distance matrix is created from BOLD time series data. (2) Next, data are split into training and held-out testing data sets. MDS is used to compress the distance matrix of the training data to a lower dimensional space. (3) Multiple regression with the MDS coordinates as the independent variables and the behavioral variable of interest as the dependent variable is used to generate a model of brain-behavior associations. (4) The model is then used in held-out test data to test predictive accuracy (i.e., model validation). (5) Following model validation, feature importance is determined via mapping edges to model predictions within the training data. Edges are then mapped back to neuroanatomy.

As shown in **Figure 4**, use of the Wasserstein metric consistently outperformed the other distance metrics, CPM and KRR in predicting IQ and this was significant after controlling for multiple comparisons (*t* (8) = 4.42, *p* = .002 vs. angular distance; *t* (8) = 4.08, *p* = .004 vs. Euclidean distance; *t* (8) = 3.78, *p* = .005 vs. Manhattan distance; *t* (8) = 4.27, *p* = .003 vs. CPM; *t* (8) = 10.89, *p* < .001 vs. KRR). Practically, this corresponded to mean increases in explained variance over CPM and KRR (i.e., *r*^2^) of ∼65-75% (CPM mean : 0.257; KRR mean *r*: 0.250; CRAM-Wasserstein mean *r*: 0.331, also see Figure S2 and Table S1). Similar improvements were observed across other common performance metrics (e.g., Spearman’s ρ, root mean squared error, coefficient of determination, Tables S2-4), as well as for prediction of IQ in an independent cohort of youth in the Philadelphia Neurodevelopmental Cohort [28] (N’s=916-1387, CPM mean *r*: 0.262; KRR mean *r*: 0.375; CRAM-Wasserstein mean *r*: 0.401, Table S5). Finally, we tested whether the improvements afforded by the Wasserstein in predicting IQ also extended to other phenotypes. Wasserstein yielded improved performance on average relative to CPM and KRR in predicting additional other HCP phenotypes, such as openness-to-experience (*t* (62) = 6.83, *p* < .001 vs. CPM; *t* (62) = 7.42, *p* < .001 vs. KRR, Table S6). Together, these results indicate that considering functional connectivity matrices as representations of their associated Gaussian distributions enables retention of features that are relevant to individual differences in phenotypic expressions, such as IQ.

**Figure 4.**
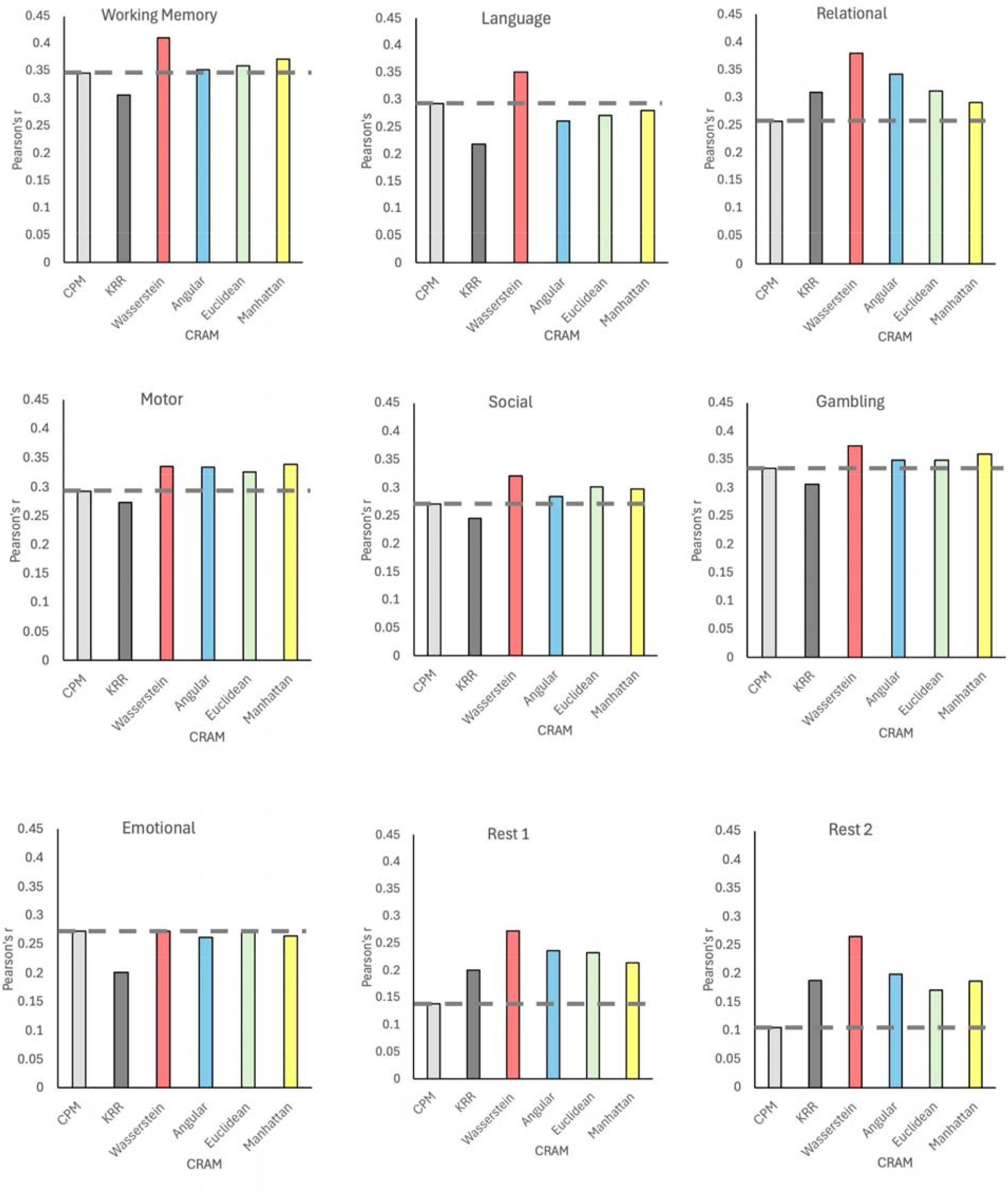
**Predictive accuracy of CPM, KRR and CRAM using different distance metrics** in predicting fluid intelligence across the seven tasks and two resting state runs from the HCP (N=515). CRAM models using the Wasserstein metric showed the highest prediction accuracy across all tasks and resting state. CPM=connectome-based predictive modeling, KRR=kernel ridge regression

### Comparison of neural networks identified using different methods and metrics

In addition to generating predictions, a fundamental goal of brain-behavior modeling is the identification of the specific brain features that drive model performance, thereby providing neurobiological insight [10]. Next, we therefore compared brain network features identified via the different modeling approaches tested above. We hypothesized that the different approaches would identify networks involving similar macroscale network-level features (reflecting that the neural substrates of IQ are biologically grounded and not wholly method- or metric-dependent) but would nonetheless differ in feature selection at the microscale, individual edge-level (reflecting the differential sensitivity of these approaches to different aspects of the matrix space).

As a benchmark for similarity, we first compared features between different iterations of 10-fold cross-validations conducted using the same method, with connections grouped by canonical networks. The stability of masks under separate runs of CRAM is comparable to the stability of masks generated by CPM. For instance, when grouping edges by canonical networks, for the gambling task in HCP, the average correlations between different positive masks generated by CRAM (CRAM-Angular *r* = 0.954, CRAM-Euclidean *r* = 0.950, CRAM-Manhattan *r =* 0.954, CRAM-Wasserstein *r =* 0.975) are comparable to the correlation between different positive masks generated by CPM (*r =* 0.947).

As shown in **Figure 5a**, the average correlation between identified network features (shown for both positive and negative predictive networks) in two separate iterations of the same method (i.e., the stability of identified features) is typically ∼.96 (range = .95-.98), whereas the mean correlation between network features from separate models from CRAM-Wasserstein vs. different methods ranged from .86-.95, indicating that features identified using the different methods and distance metrics are similar but not identical. Collectively, this suggests that brain-behavior modeling approaches reliably identify specific neurobiological features subserving individual differences in phenotypic expressions in a manner that is largely robust to different methods and metrics, and that considering matrices as representations of their associated Gaussian distributions enables retention of connectivity features that are not always captured by vector-based approaches.

**Figure 5.**
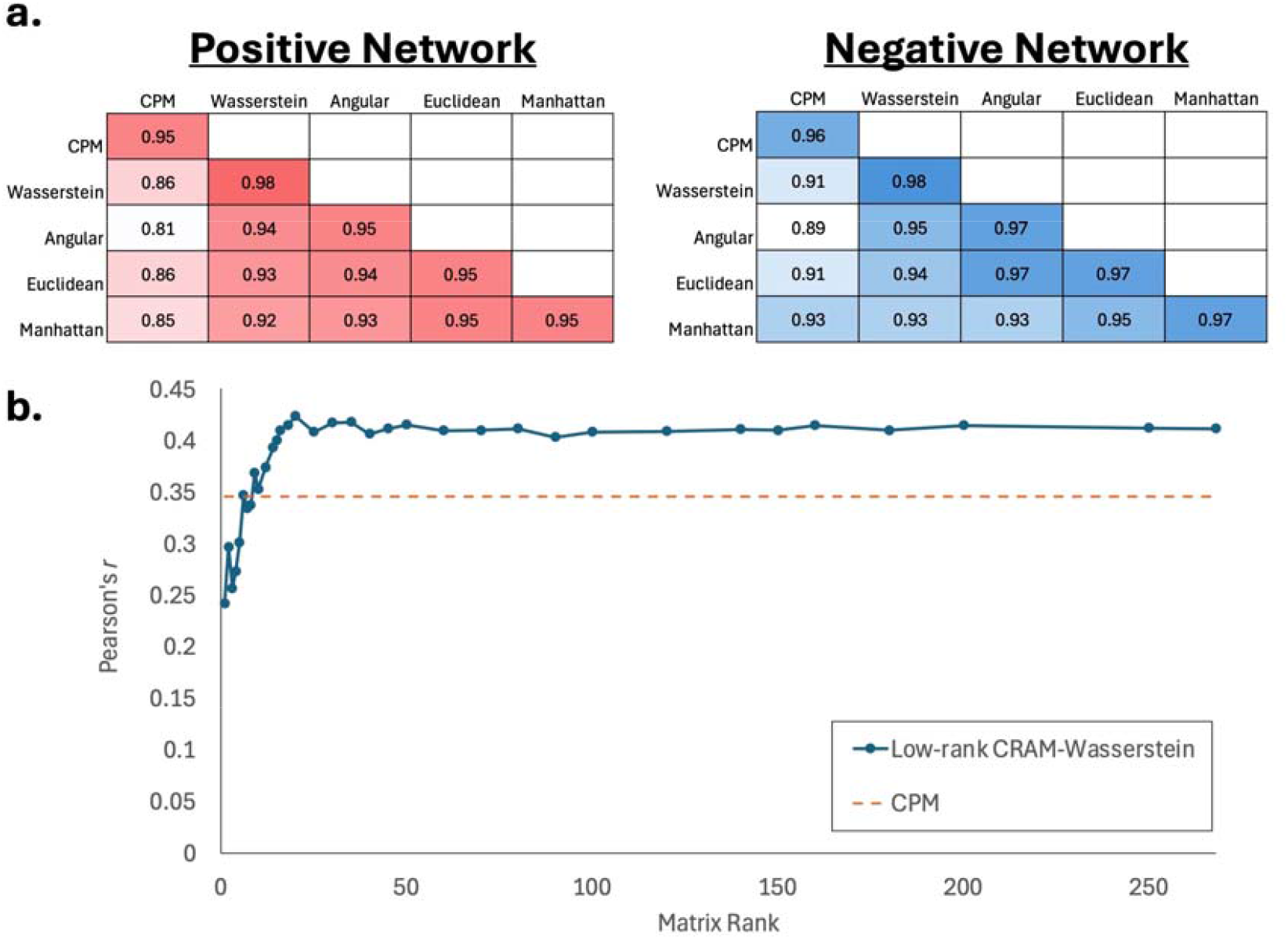
Network feature similarity across methods. A. The correlation between identified network features in two separate model runs. The correlation between network features from separate runs using different methods is lower than those using the same method, indicating that features identified using the different methods and distance metrics are similar but not identical. B. The prediction accuracy using successively smaller eigenvectors in CRAM-Wasserstein versus CPM. Adding in additional components changes model performance, with an initial small decrease followed by a steady increase which plateaus to the level of CRAM-Wasserstein once 15 components are incorporated

### Implications for understanding the human connectome

We next asked what features of the matrix space might be contributing to the improvements in predictive accuracy afforded by the Wasserstein. We first hypothesized that performance increases were due to the ability of this metric to capture additional information regarding third-order interactions between brain regions (which are not considered in standard vector-based approaches which only consider pairwise interactions). To test this, we extended the pairwise 268×268 matrices from the HCP by creating 268×268×68 tensors of the normalized coskewness between triads of nodes for all matrices in the dataset (see Methods). Normalized coskewness, like correlation, takes an average of the product of deviations from the mean and divides it by the product of standard deviations of the three variables. Then, as in CPM, we correlated each element in the 3D coskewness tensor with an outcome variable (fluid intelligence as above) to identify features in training data and to build a linear predictive model. This third-order version of CPM (hereafter referred to as C3PM), performed significantly worse than standard CPM at prediction of intelligence (*r*_*C3PM*_ = 0.264, *r*_*CPM*_ = 0.344, *t* (38) = 17.92, *p* < .001). As a further test, we combine features identified from CPM and C3PM in a cross-validated multiple regression framework. Results indicated that consideration of local third-order statistical interactions in the matrix space does not increase—but in fact decreases—predictive accuracy (*r _multiple_* = 0.269, *r*_*CPM*_ = 0.344, *t* (38) = 9.91, *p* < .001). Therefore, it is highly unlikely that the Wasserstein’s improved performance is attributable to consideration of these triadic relationships. By extension, this suggests that such third-order statistical interactions between nodes do not encode for meaningful individual differences above and beyond what is captured by standard pairwise nodal interactions.

Additionally, we used these coskewness tensors in place of standard FC matrices to calculate ID rates. Specifically, we used the vectorized coskewness tensors for each participant per scan, calculated pairwise angular distances, and used those distances as described in the previous section on ID rate calculations. Results were poor: the average ID rate between all pairs of scans in the HCP data set was 6.9% (vs. 93.6% when using the Wasserstein distance), and for several scan pairs the ID rate was no better than chance. We interpret this finding as evidence that three-way interactions among BOLD time series do not provide sufficient information to explain the improvements in ID rate due to incorporating the Wasserstein metric.

We next hypothesized that the Wasserstein metric’s ability to retain information about both local (i.e., as in pairwise nodal interactions) *and* global properties of connectivity matrices might be driving the observed improvements in performance. To test this, we sought to determine the predictive accuracy of models built using only global summary metrics—as would be equivalent to the principal component or eigenvector of the entire matrix—either alone or in combination with successively smaller components, relative to models built using the entire distribution (as in the Wasserstein). In other words, we reasoned that if both local and global properties of the matrix space were contributing to model performance, then removing all information from the matrices apart from the principal component/eigenvector should maintain most of the signal, *and* re-introducing information from a small number of the next largest eigenvectors should generate the full improvement seen from the Wasserstein metric.

To test this second hypothesis, we generated low-rank approximations of each connectivity matrix from the HCP. We decomposed the matrices into their eigenvector/value pairs, zeroed out all eigenvalues except for the first *k*, and then reconstructed the matrices. For selected values of *k* from 1 to 268, we performed CRAM with the Wasserstein on the reconstructed working memory task connectivity matrices and compared the result against CPM (which only considers local statistical interactions).

As shown in **Figure 5b**, models built using only the principal component (i.e., the first principal eigenvector) of each matrix contained enough information for a predictive r of .24 (which is lower in performance than standard CPM). Adding additional successively smaller components further impacted model performance, with an initial small decrease followed by a steady increase which surpassed CPM after 6 components were added and which plateaued to the level of the full Wasserstein model once 18 components were incorporated. Thus, while CPM has better predictive accuracy than a single principal component model, the addition of a small number of other components adds enough information to outperform CPM. Taken together, this suggests that one factor driving performance of our distribution-based approach is the method’s increased consideration of global relationships, such as covariance in connectivity across all brain regions. At the same time, the improved performance of CPM (which only considers local interactions) relative to the first principal component (which only considers global properties) suggests that both local *and* global properties are important for model accuracy and thus encode for meaningful individual differences in IQ.

Furthermore, we used these low-rank FC matrix approximations to calculate ID rates based on the Wasserstein distance. Specifically, for each rank *k* approximation, we computed the average of all ID rates between scan pairs based on the Wasserstein distance between the reduced rank FC matrix approximations. As shown in Figure XXX, a 50% ID rate required inclusion of six components per FC matrix. Surpassing the equivalent of the Angular distance ID rate (68.6%) required retaining ten components. To reach the ID rate achieved with the Wasserstein distance on the unaltered FC matrices (93.6%) required keeping 40 components per matrix. As noted above, full CRAM predictive accuracy was obtained after adding only 18 components; we take this as evidence that more aspects of the FC matrix contribute to ID rate improvements as compared with predictive accuracy improvements. The exact reasons why this occurs are unclear, and examining why this is the case is beyond the scope of this paper.

## Discussion

Common methods for analyzing the human connectome often focus on pairwise relationships between two regions or nodes, referred to as edges, as summarized in a connectivity matrix. However, edges within a connectivity matrix are not independent [18] - they are part of an interconnected system. Using neuroimaging data from hundreds of individuals, we show that adopting a global, geometrically grounded measure of similarity - the Wasserstein distance metric - outperforms existing popular methods in distinguishing between individuals (aka connectome ‘fingerprinting’) [1] and in building robust brain-behavior predictive models of fluid intelligence. In doing so, we introduce a novel flexible regression pipeline with built-in cross-validation, connectome regression in any metric (CRAM, https://github.com/misterriley/CRAM), which we developed to test differences in predictive model accuracy across different metrics. We demonstrate that, when paired with the Wasserstein, this approach outperforms both connectome-based predictive modeling (CPM) and kernel ridge regression (KRR) methods of brain-behavior modeling, resulting on average in increases of 65-75% in explained variance for models of IQ.

Considering the entire matrix space as a distribution is more computationally burdensome than treating each value independently. To address this, CRAM incorporates dimensionality reduction using multidimensional scaling (note that this transformation is learned from the training data and applied out-of-sample to the testing data, preventing leakage). One concern when conducting dimensionality reduction is that recovery of meaningful network features may be more challenging, thereby reducing direct mapping of model findings to brain anatomy — a central goal of brain-behavior modeling [10]. In addition, networks identified using connectome-wide predictive modeling approaches are typically complex (composed of many edges connecting distributed brain regions) [1, 6, 29]. While this likely reflects the true nature of networks supporting complex processes such as cognition [1, 30] and clinical change [6, 31], the inherently multivariate nature of these approaches nonetheless risks identification of spurious connections, even with stringent cross-validation [16, 17, 32]. However, our analyses demonstrated that, not only are features selected across different iterations of the same method and metric relatively consistent, but that feature overlap at the level of macroscale brain features – i.e., number of edges identified from a given canonical neural network - is also high across different methods and metrics, even those conducted with and without multidimensional scaling. Despite this, using connectivity matrices computed from both resting-state and all seven tasks and acquisitions, the distribution-based Wasserstein metric consistently outperformed vector-based distance metrics, such as Euclidean and Angular distances. This suggests that the improvements afforded by the Wasserstein are not solely attributable to CRAM’s use of dimensionality reduction but are likely related to the use of a distribution-based metric itself.

The Wasserstein metric quantifies distance over the space of probability distributions. Practically, this means that computation of the Wasserstein distance both incorporates and depends upon the geometry of the curved manifold of covariances matrices. In contrast, vector-based metrics are only informed by the flat structure of the Euclidean distance that surrounds the smaller space of covariance matrices. Thus, in choosing a distance metric that retains information about the underlying structure of the matrix, we leverage both local, edge-level and global, distribution-level features during predictive model creation. This approach thereby enables brain-behavior modeling with retention of two key features which are not considered in standard pairwise vector-based approaches: (i) higher order statistical associations, such as third-order interactions between triads of local brain regions and (ii) global statistical properties, such as those corresponding to the dominant mode of connectivity co-variation across all brain regions.

To test which of the above properties might be driving the increase in model performance afforded by the Wasserstein, we conducted a series of post-hoc analyses: First we tested whether performance increases were due to the ability of the Wasserstein metric to capture information about local, third-order associations between brain regions by comparing predictive ability of models generated using 268×268×268 tensors vs 268×268 matrices. Results indicated that consideration of such local, third-order interactions actually *decreased* model performance relative to models only considering pairwise relationships. We next considered whether the metric’s consideration of global properties was driving improvements in performance via testing the predictive ability of global metrics, in this case principal eigenvectors, alone and in combination with local properties (as defined by increasingly smaller eigenvectors). Taken together with results of CPM analyses (which only consider local interactions yet outperformed analyses using solely the first eigenvector), this suggests that both local pairwise *and* global properties meaningfully encode for individual differences in measurable phenotypes such as IQ, and should therefore be considered in brain-behavior modeling approaches. Critically, these individual differences in the geometry of the curved manifold of connectivity matrices (e.g., difference in space between primary modes of brain covariation, or the difference between eigenvectors) are not considered in standard brain-behavior modeling approaches.

Several limitations should be noted, including our focus on a single distribution-based metric — the Wasserstein. While we considered other metrics for comparing probability distributions (e.g., Fisher-Rao, Jensen-Shannon divergence, Hellinger distance), these metrics are more sensitive to measurement noise in cases where the ratio between the number of data points and the number of data dimensions is relatively low [33, 34], as may be the case when creating functional connectivity matrices (details in Supplement). In addition, metrics such as Jensen-Shannon divergence and Hellinger distance are functions of the ratio of probabilities between two distributions resulting in increased sensitivity to the tails of a given distribution (which are by definition more variable and less representative of the majority of the distribution) [35, 36].

Further, other candidate distance metrics, such as Mahalanobis distance and Log-Determinant divergence, are functions of the similarity between inverse covariance matrices, which have increased sensitivity to measurement noise [33]. Additional limitations include our primary focus on a single behavioral variable – fluid intelligence – from a single dataset – the Human Connectome Project (HCP). While we also demonstrated the utility of our approach to predict a small number of additional phenotypes from HCP, and to also predict IQ using data from the Philadelphia Neurodevelopmental Cohort, further testing of the robustness of our approach to other datasets – including those acquired without harmonized acquisition parameters and focusing on prediction of more complex phenotypes such as future clinical outcomes – is nonetheless warranted.

These results indicate that the human connectome contains multiple sources of information that may be used to reliably distinguish between individuals and to generate individual-level predictions of fluid intelligence and other phenotypes. We demonstrate that, in addition to individual differences in connectivity between pairs of regions, individual differences in more global properties, such as those defined by the overarching geometry of an individual’s connectivity space, carry critical information about the relative uniqueness of an individual’s functional brain organization.

## Acknowledgements

This work received support from the following sources: National Institute on Drug Abuse T32DA022975 (SR), National Institute of Mental Health grant R01MH120080 (AH), National Institute of Biomedical Imaging and Bioengineering grant R01EB034720 (YZ).

## Data and code availability

The scripts for running the analyses conducted in the study are available at https://github.com/YaleYipLab/CRAM.

## Author contributions

The overall study concept was created by SR in consultation with SWY, AC, TC and XS. Code was written by SR. Analyses were conducted by SR and SC. Manuscript writing was conducted by SR, SC and SWY. All authors provided critical feedback on this project and have approved the final manuscript.

## Ethics

This study used data from the Human Connectome Project. All subjects gave signed informed consent in accordance with the protocol approved by the institutional review board associated with the HCP.

