## Supplemental information for "Is the whole more than the sum of its parts? Considering global and local features of the connectome improves prediction of individuals and phenotypes"

**Supplement**

*Metrics*

*Norms as metrics*

*Correlation as compared to angle between vectors*

*Pre-processing and matrix generation*

*MDS dimensionality*

*Third-order Connectome-based Predictive Modeling (C3PM)*

*Low-rank CRAM-Wasserstein*

*Differences in identified networks by significance threshold*

*Virtual lesioning*

*Other metrics not considered*

*Table S1. Predictive accuracy by method and metric, as measured by Pearson’s* $r$

*Table S2. Predictive accuracy by method and metric, as measured by Spearman’s ρ*

*Table S3. Predictive accuracy by method and metric, as measured by coefficient of determination*

*Table S4. Predictive accuracy by method and metric, as measured by root mean squared error*

*Table S5. Clustering coefficients of identified networks by method and metric.*

*Table S6. Predictive correlation using CRAM-Wasserstein vs. MDS dimension*

*Figure S1. Average ID rate for all possible pair-wise combinations of 9 scans*

*Figure S2. Average predictive accuracy for CPM, KRR and CRAM in predicting fluid intelligence across all 9 scans*

### Metrics

A *metric* is a function that formalizes the notion of distance between two objects. To be a metric, a non-negative function 𝑑 over a space 𝑋 must satisfy three properties for all $x,y,z\in X$:

1) $d\left( x,y \right)=0$ if and only if $x=y$
2) $d\left( x,y \right)=d\left( y,x \right)$
3) $d\left( x,y \right)\leq d\left( x,z \right)+d\left( z,y \right)$

Property 1 ensures that distinct points must have some positive distance between them. Property 2 requires that distances are the same going each direction. The third property, known as the triangle inequality, indicates that it is not possible to add an intermediate destination while decreasing the travel distance. Any function that satisfies all these properties is a metric. The space $X$ equipped with the metric $d$ is then known as a *metric space*.

Some functions (such as angular distance, discussed below) satisfy properties 2 and 3, but not property 1. These functions are often called *semi-metrics*.

### Norms as metrics

A *norm* is a nonnegative function *f* that formalizes the notion of length over a vector space $V$. Like a metric, a norm must satisfy three properties. For all $x,y\in V$:

1) $f\left( x \right)=0$ if and only if $x=0$
2) $f\left( ax \right)=\left| a \right|*f\left( x \right)$ for any scalar $a$
3) $f\left( x+y \right)\leq f\left( x \right)+f\left( y \right)$

Properties 1 and 3 are directly analogous to properties of metrics. Property 2 requires that scaling a vector also scales the length of that vector.

The vector space $V$ equipped with the norm $f$ creates a *normed space*. Due to these properties, a norm can *induce* a metric. Specifically, if we define $d\left( a,b \right)=f\left( a-b \right)$, then $d$ satisfies the properties of a metric. The most well-known norm is the Euclidean norm, commonly written as $f\left( x \right)=\left\| x \right\|$ or $\left\| x \right\|_{2}$, which can be computed using the Pythagorean theorem. If $x$ is an $n$-dimensional vector with components $\left( x_{1},x_{2},\ldots,x_{n} \right)$, then:

$$\left\| x \right\|_{2}=\left( \sum_{i=1}^{n} x_{i}^{2} \right)^{\frac{1}{2}}$$

. This formula can be generalized to the class of 𝑝-norms by replacing the 2s with a value $p\geq1$:

$$\left\| x \right\|_{p}=\left( \sum_{i=1}^{n} \left| x \right|_{i}^{p} \right)^{\frac{1}{p}}$$

. Besides the Euclidean norm ($p = 2$), another commonly used norm is the Manhattan norm ($p=1$), named because paths between locations in Manhattan are constrained to a rectangular grid. These two norms induce the Euclidean and Manhattan distances used in this paper.

Manhattan distance:

$$d\left( x,y \right)=\sum_{i=1}^{n} \left| x_{i}-y_{i} \right|$$

Euclidean distance:

$$d\left( x,y \right)=\left( \sum_{i=1}^{n} \left( x_{i}-y_{i} \right)^{2} \right)^{\frac{1}{2}}$$

### Correlation as compared to angle between vectors

Here we describe the relationship between angle and correlation of two vectors. Consider $x$ and $y$ with respective means $\bar{x}$ and $\bar{y}$, each of which are vectors of length $n$. The Pearson’s correlation ($r$) between these vectors is their covariance divided by the product of their standard deviations. After some simplification:

$$r_{xy}=\frac{Cov\left( x,y \right)}{\sigma_{x}\sigma_{y}}=\frac{\sum_{i=1}^{n} \left( x_{i}-\bar{x} \right)\left( y_{i}-\bar{y} \right)}{\sqrt{\sum_{i=1}^{n} \left( x_{i}-\bar{x} \right)^{2}\sum_{i=1}^{n} \left( y_{i}-\bar{y} \right)^{2}}}$$

. If $x$ and $y$ are centered vectors (i.e., $\bar{x}=\bar{y}=0$) then this equation further simplifies to

$$r_{xy}=\frac{\sum_{i=1}^{n} x_{i}y_{i}}{\sqrt{\sum_{i=1}^{n} {x_{i}}^{2}\sum_{i=1}^{n} {y_{i}}^{2}}}$$

. Separately, to find the angle $\theta$ between the vectors we use the cosine angle formula:

$$\cos\left( \theta\right)=\frac{x\cdot y}{\left\| x \right\|\left\| y \right\|}$$

. The value of $x\cdot y$ (the dot product of the vectors) is $\sum_{i=1}^{n} x_{i}y_{i}$, and $\left\| x \right\|$ (the Euclidean norm of the vector) is $\sqrt{\sum_{i=1}^{n} {x_{i}}^{2}}$. Thus:

$$\cos\left( \theta\right)=\frac{\sum_{i=1}^{n} x_{i}y_{i}}{\sqrt{\sum_{i=1}^{n} {x_{i}}^{2}\sum_{i=1}^{n} {y_{i}}^{2}}}$$

. This coincides with the formula for correlation between centered vectors, so we conclude that

$$r_{xy}=\cos\left( \theta\right)$$

when $x$ and $y$ both have mean 0. We see from this formula that correlation is closely related to a function of the angle between vectors when the mean of those vectors is equal to or approximately zero.

Note that angular distance violates property 1 of a metric. When scaling a vector, the direction does not change; thus, angular distance does not change with length. Since the angle between 𝑥 and itself is 0, so must the angle from 𝑥 to 𝑎𝑥 be 0 for any scalar $a>0$. This violates the condition that different vectors must always have a positive distance between them.

### Pre-processing and matrix generation

Neuroimaging data were obtained from the Human Connectome Project (HCP) 900 Subjects release[1]. From this data set, we used behavioral and functional imaging data. Details on pre-processing and matrix generation have been reported previously[2]. Briefly, our analyses focused on participants who completed all nine fMRI runs (seven task-based and two resting-state), exhibited minimal motion artifacts—defined by mean frame-to-frame displacement below 0.1 mm and maximum displacement below 0.15 mm—and had available general cognitive ability (gF) scores (final sample: n = 515; 241 males; age range: 22–36+). These stringent motion criteria, consistent with HCP preprocessing protocols, were implemented to minimize motion-related confounds in functional connectivity estimates. Notably, after applying these thresholds, motion was not significantly correlated with gF in the majority of scanning conditions (all *P* > 0.05, Bonferroni corrected) except the social task, right-left (RL) phase encoding run (*r_s_* = −0.16 (*P* = 0.00017)), the relational task, left-right (LR) phase encoding run (*r_s_* = −0.15 (*P* = 0.0008)), and the emotion task, RL phase encoding run (*r_s_* = −0.14 (*P* = 0.0017)).

Analyses involving HCP data were classified as exempt from IRB review by the Yale Human Investigation Committee. All HCP imaging data were processed using the minimal preprocessing pipeline, which includes artifact removal, motion correction, and alignment to a standardized template. Additional preprocessing was carried out using BioImage Suite and involved standard procedures such as regressing out motion-related signal components, mean signals from white matter, cerebrospinal fluid, and gray matter, as well as linear trend removal and low-pass filtering. For each subject, average frame-to-frame displacement was computed across both LR and RL phase encoding runs, resulting in nine motion estimates per individual; these values were subsequently used for participant exclusion and motion-related analyses.

Functional connectivity was assessed using the 268-node Shen atlas, which was generated from an independent dataset using group-wise spectral clustering. After parcellating each individual’s preprocessed data into these functionally defined nodes, pairwise correlations between node time courses were calculated and transformed using Fisher’s r-to-z transformation, yielding nine 268 × 268 connectivity matrices for each HCP participant. For the PNC dataset, three such matrices were generated per subject (one per fMRI run). In the case of the HCP data, matrices were computed separately for LR and RL runs and then averaged within each condition. Nodes with inadequate spatial coverage—typically due to limited scan volume—were excluded from all subjects in a conservative fashion. Specifically, nine nodes were found to have insufficient coverage in the HCP data and were excluded from all subsequent analyses.

### *Multidimensional scaling (MDS) dimensionality reduction*

MDS is a dimensionality reduction technique that minimizes the *stress* caused by moving a set of points from a space equipped with any metric-like dissimilarity measure into a space equipped with the Euclidean norm. To accomplish this transformation, MDS converts any distance structure for a given set of points into a ‘flat’ space while respecting the original distances [3]. Conceptually, MDS can be visualized as acting on a set of interconnected balls bound together by springs, where the relaxed length of the spring connecting points $p_{i}$ and $p_{j}$ equals the entry at position $i,j$ in the dissimilarity matrix $D$. MDS projects these balls and springs into a space of dimension $k$, specified by the user, where the resulting arrangement finds the lowest energy configuration for the set of springs. Let $y_{i}$ be the output of $p_{i}$. The stress $S$ of the transformation is defined as:

$$S=\sum_{i} \sum_{j>i} \left( D_{ij}-\left\| y_{i}-y_{j} \right\| \right)^{2}$$

Applying MDS to the matrix $D$ yields a set of points that minimizes $S$.

Linear regression applied to the outputs of MDS reflects the dissimilarity measure that was chosen in the original space. Since the dissimilarity matrix only needs to be symmetric, it allows for an insertion point for any distance-like measurement between pairs of points, including angles between vectors, norm-based distances, and matrix or probability distribution distance measures, such as the Wasserstein metric.

CRAM, which uses MDS for dimensionality reduction, requires specification of the number of dimensions $d$ in which to embed data. We chose to use a common rule of thumb that specifies $d$ based on the eigenvalues of the transformation matrix. Specifically, we retain all dimensions where the associated eigenvalue is at least as large as the average eigenvalue across all dimensions. Some data require only a few dimensions for an embedding that maintains distances through the transformation. Such a data set would generate a transformation matrix with a few large eigenvalues and many small ones; thus $d$ would be small.

We tested whether CRAM-Wasserstein’s predictive accuracy, averaged over all nine HCP brain states, changed significantly with the rule of thumb we chose. For values of $t$ from 50% to 150%, we multiplied the eigenvalue threshold by $t$ before selecting $d$. That is, if $t$ was 50%, we retained all dimensions where the eigenvalue was at least 50% of the mean of all the eigenvalues of the transformation matrix. We then ran CRAM-Wasserstein using the computed value of $d$ and calculated the average Pearson’s $r$ value for predictions across 100 runs.

Results from this analysis are reported in Table S1 and Figure S1. Average predictive accuracy across the nine brain states ranged from $r=.305$ for $t=50\%$ to $r=.334$ for $t=120\%$. As such, the predictive accuracy of CRAM-Wasserstein under the $t = 100\%$ rule of thumb was very slightly less than the maximum for the tested values. We conclude the chosen rule for selection of $d$ is not a critical factor in the performance of CRAM.

### Third-order Connectome-based Predictive Modeling (C3PM)

FC matrices are composed of correlations between timeseries, and as such, the entries in FC matrices are not independent from each other. That is, if the correlation between nodes $x$ and $y$ is known and so is the correlation between nodes $y$ and $z$, this limits the range of valid correlation coefficients between nodes $x$ and $z$. These types of constraints hold for every set of three or more timeseries.

The result of these constraints is that FC matrices, like all correlation matrices, must be positive semi-definite (PSD). PSD matrices have eigenvalues (defined in the next section) that are non-negative. The result of the PSD constraint is that the space of valid correlation matrices among larger sets of nodes shrinks exponentially as a fraction of all matrices that would otherwise look like correlation matrices (i.e., those that are symmetric with diagonal equal to 1 with all entries bounded by 1 and -1). The geometry that is induced by these constraints has been leveraged before in the study of functional connectivity [4].

CPM and other commonly used methods for analyzing functional connectivity do not employ information about interactions among more than two nodes at a time. Since the full correlation structure is used to calculate the Wasserstein metric, which requires valid covariance matrices as inputs (i.e., the Wasserstein distance calculation yields a complex number if an input is not a covariance matrix), it follows that the Wasserstein metric is sensitive to violations of the constraints on FC matrices. This sensitivity is because the Wasserstein metric measures the cost of transforming one covariance matrix into another while respecting the geometry of the space of PSD matrices. In contrast, many other distance metrics (e.g., Euclidean distance) and methods (e.g., CPM) treat the entries of the matrix as independent, ignoring the PSD constraint and the resulting higher-order dependencies among correlations. By respecting the PSD constraint, the Wasserstein metric implicitly captures the influence of three-way and higher-order interactions among time series, as these interactions are necessary to maintain the validity of the correlation structure.

We therefore hypothesized that the improvements in the CRAM-Wasserstein method against other methods and metrics were due to the ability of the Wasserstein metric to capture the influence of three-way and higher-order interactions among timeseries. If true, this would imply that extending CPM to include information about third-order interactions should improve predictions beyond what is found in standard CPM. To test this hypothesis, we created a modified version of CPM that takes as its input a three-dimensional tensor of three-way interactions among time series. We call this method third-order CPM, or C3PM for short.

FC matrices are composed of pairwise correlations $r$ between the BOLD timeseries for each pair of nodes. Let $\bar{x}$ and $\bar{y}$ be the mean of the timeseries for nodes $x$ and $y$, respectively. The formula for correlation between $x$ and $y$ is

$$r_{xy}=\frac{\sum_{i=1}^{n} \left( x_{i}-\bar{x} \right)\left( y_{i}-\bar{y} \right)}{\sqrt{\sum_{i=1}^{n} \left( x_{i}-\bar{x} \right)^{2}\sum_{i=1}^{n} \left( y_{i}-\bar{y} \right)^{2}}}$$

. An analogous measure for a set of three nodes $x$, $y$, and $z$ is

$$s_{xyz}=\frac{\sum_{i=1}^{n} \left( x_{i}-\bar{x} \right)\left( y_{i}-\bar{y} \right)\left( z_{i}-\bar{z} \right)}{\sqrt{\sum_{i=1}^{n} \left( x_{i}-\bar{x} \right)^{2}\sum_{i=1}^{n} \left( y_{i}-\bar{y} \right)^{2}\sum_{i=1}^{n} \left( z_{i}-\bar{z} \right)^{2}}}$$

. This measure is the *normalized coskewness* of the three timeseries. Intuitively, $s_{xyz}$ is a measure of how the correlation between $x$ and $y$ changes as a function of $z$.

For each scan we create a 3D tensor of normalized coskewness values, with one entry for every combination of three nodes in the atlas. Then the tensors are vectorized, and the steps of standard CPM are run on these vectors. Each location in the vectors is checked for a significant correlation with a behavioral measure at the $p < .05$ level, producing a mask of 1, 0, and -1 values, which indicate positive, non-significant, and negative correlations, respectively. In held out data, summing over the product of a vector times the mask creates a summary connectivity value. Cross-validation produces a summary connectivity value for each data point. This summary value is assessed for predictive power by seeing how well it correlates with the behavioral variable across the full data set. Additionally, this variable can be paired with the summary variable from standard CPM and used for multiple regression.

We test CPM vs. C3PM in the HCP working memory task averaged over 20 runs. Results from these tests are shown in figure XX. C3PM underperforms standard CPM by a significant margin ($r_{C3PM} = 0.264$, $r_{CPM}=0.344$, $t(38) = 17.92$, $p < .001$) when its summary value is used alone. When paired with the CPM summary value for multiple regression, the result is still worse than CPM ($r_{multiple}=0.269$, $r_{CPM}=0.344$, $t(38)=9.91$, $p<.001$). This finding goes against the hypothesis that information from three-way interactions improves predictive accuracy. This negative result might be due to the increased size of the input vector (~32 million entries in C3PM vs. ~35 thousand for CPM). This explosion in terms might point to the need for a different $p$-value cutoff to reduce the number of spurious non-zero entries in the mask. Perhaps using a more stringent *p*-value for thresholding would yield better performance. Alternatively, other methods for paring down which sets of $s_{xyz}$ values to look at might improve performance. We did not examine these possibilities but instead chose to examine the relationship between global associations and predictions.

Next, we used angular distances between vectorized coskewness tensors to see how well this measure of distance performs when calculating ID rate. For a given participant and a given brain state, we take the set of $s_{xyz}$ values for $x\geq y\geq z$ and stack them into a vector of size 32080940. We then calculate angular distance between pairs of coskewness vectors and compare distances between scans using the ID rate method. Results were substantially worse than ID rates with other distance measures. The average ID rate between pairs of scans in the HCP dataset was 6.9%, compared with 93.6% for the Wasserstein distance and 68.6% for the angular distance on FC matrices. We interpret this low rate of correct identifications as evidence that the information contained in the three-way interactions between node timeseries is not sufficient to generate the large improvements found when calculating ID rates via the Wasserstein distance.

### Low-rank CRAM-Wasserstein

Given the lack of support for the previous hypothesis, we sought to understand whether global coactivation patterns, as captured by low-rank approximations to FC matrices, could explain the predictive power of CRAM-Wasserstein. These low-rank approximations were generated by retaining only a subset of eigenvalue/vector pairs and zeroing out all others, then running CRAM using the Wasserstein distance between the reduced rank matrices.

An *eigenvector* $v$ of a square matrix $A$ is a vector that, when multiplied by $A$, yields a vector that only differs from $v$ by a scalar multiplier $\lambda$. This scalar multiplier is known as the *eigenvalue* associated with $v$:

$$Av= \lambda v$$

*.*

*Eigendecomposition* is a method for obtaining and collating all eigenvectors and eigenvalues of a matrix. Let $Q$ be a square matrix where column $i$ is the $i$th eigenvector of $A$ and let $\Lambda$ be a diagonal matrix where the value at $\Lambda_{ii}$ is the $i$th eigenvalue of $A$. The eigendecomposition of A yields $Q$ and $\Lambda$ such that

$$A= Q\Lambda Q^{-1}$$

. When A is symmetric, the eigendecomposition exists. When the eigendecomposition exists, the number of non-zero entries in $\Lambda$ is known as the *rank* of the matrix. FC matrices are symmetric and PSD, meaning that there are no eigenvalues less than zero.

The eigendecomposition can be used to construct an approximation of $A$ by replacing some entries in $\Lambda$ with zero, yielding a reduced rank matrix that maintains the PSD constraint. Let $\Lambda_{g}$ be the result when all but the largest $g$ values in $\Lambda$ are replaced by zero. We define the rank-$g$ approximation to $A$ as

$$A_{g}= Q\Lambda_{g}Q^{-1}$$

. Retaining the largest eigenvalues ensures that these low rank approximations keep the strongest global coactivation signals; discarding the smaller eigenvalues removes measurement noise and weaker interactions.

Using this formula, we compute the rank-$g$ approximation for all FC matrices. We then use CRAM with the Wasserstein metric applied to the set of rank-$g$ approximations for select values between $g=1$ to $g=268$, predicting fluid intelligence from the working memory scan of HCP. Results can be seen in figure XX. When $g=1$, the correlation between predictions and actual values is $r=.24$, indicating that a large fraction of the predictive power of the scan lies in the principal eigenvectors of the matrices. The predictive accuracy increases rapidly as eigenvectors are added back into the matrices. When $g = 6$, this method outperforms CPM, and when $g = 18$, the method achieves the same results as CRAM-Wasserstein. This outcome shows that only a small number of eigenvectors of the FC matrices are needed for the increases in predictive accuracy of CRAM-Wasserstein vs. other metrics and vs. CPM, and that any extra information contained in the smaller eigenvalues is either irrelevant to this prediction or contains too much noise to be useful.

Additionally, we tested how well we were able to identify participants between scans when using these reduced rank FC matrices. For selected values of *g* from 1 through 268, we computed the average ID rate between all pairs of HCP brain states using the Wasserstein distance on the set of rank *g* matrices. Results are shown in figure XXX. When $g = 1$, ID rate is 13.7%, but climbs steadily with increases in *g*. At $g = 6$, ID rate reaches 50%, and when $g = 10$, ID rate reaches the 68.6% ID rate of angular distance on the full FC matrices. To obtain the 93.6% ID rate when using the Wasserstein distance between full matrices requires $g = 40$. Surprisingly, ID rate peaks above the full matrix Wasserstein ID rate at 95.4% when $g = 50$ ($z = 10.75$, $p < .001$, two-sided two-proportion-difference z-test). This may be an indication that the smaller eigenvalues of these matrices are adding more noise to the outputs of the ID rate tests than they are adding signal. Of note is the fact that full predictive accuracy using CRAM-Wasserstein occurs at $g = 18$, while full ID rate using the Wasserstein distance requires $g = 40$. The difference in the required rank of matrices for full performance under these two tests suggests that some features of the matrices are affecting the outcomes differentially. The exact causes of this difference are unclear and beyond the scope of this paper; future research may be able to answer this question more fully.

### Differences in identified networks by significance threshold

To recover edge networks associated with task outcomes for a given model, we used post-hoc permutation testing at a two-sided significance level of $\alpha= .05$ uncorrected. That is, for a set of input matrices $X$ paired with outcome scalars $y$, we created 1000 permutations of the set of *y* values, labeled as $y_{m}$ for $m=1$ through $m=1000$. For each *m* we create a set of outcome predictions $\hat{y_{m}}$ using CRAM with inputs *X* and outcomes $y_{m}$ based on a chosen distance measure. For each edge location $i,j$ in *X*, we generate correlation $r_{ij,m}$ as the correlation between the predicted output values $\hat{y_{m}}$ and the corresponding entries $X_{ij}$ across all participants. For the true model predictions $\hat{y}$, we can also calculate corresponding values $R_{ij}$. If $R_{ij}$ falls outside of the center 95% of the values $r_{ij,m}$, then we determine that edge to be significant. Define the positive mask $M^{+}$ to be the matrix of positive edges: $M_{ij}^{+}=\left\{ \begin{aligned} 1 if R_{ij}\geq r_{ij,m} for 97.5\% of m \\ 0 otherwise \end{aligned} \right\}$. The corresponding negative mask $M^{-}$ is defined as: $M_{ij}^{-}=\left\{ \begin{aligned} 1 if R_{ij}\leq r_{ij,m} for 97.5\% of m \\ 0 otherwise \end{aligned} \right\}$.

We use the $\alpha= .05$ uncorrected significance level to maintain consistency with CPM, which also typically uses this level for significant edge detection. We sought to test how different α values affect the detected masks. First, we checked whether this threshold produced masks that were stable between runs. To test for stability, we first group the 268 nodes of the Shen atlas into the ten canonical networks identified in [5]. We then count the number of significant edges that connect nodes between each pair of canonical networks, creating a ten-by-ten matrix out of these counts. We then vectorize the lower triangle of this matrix (including the diagonal) to remove repeated entries. For two identified masks, when collapsed into these vectors, we can then calculate the correlation between the corresponding vector elements. We define stability as the average correlation between these vectorized canonical network masks produced from different runs of the same method. We can also use this analysis to compare similarity of masks between different methods, CRAM using different distance measures, and different thresholds. All tests in this section use the working memory scan from the HCP data set while predicting fluid intelligence. Results of this analysis can be found in table XXX.

Using $\alpha= .05$, the average correlation between positive masks of different CRAM-Wasserstein runs is $r=.986$. This is comparable to the average correlation between consensus CPM masks ($r=.981$), which are generated by taking only edges that are significant across all cross-validation folds for a run of CPM. When decreasing the threshold to $\alpha= .01$, the stability of CRAM-Wasserstein masks decreases to $r=.954$, and at $\alpha= .00$ the average correlation drops to $r=.883$. So, in addition to being consistent with the CPM threshold, $\alpha= .05$ produces highly stable edge masks.

Different thresholds also produce slightly different masks. The average correlation between CRAM-Wasserstein masks at $\alpha= .05$ and $\alpha= .01$ is $r = .929$, but this value drops to $r=.741$ when comparing $\alpha= .05$ to $\alpha= .00$. By comparison, the average correlation between CRAM-Wasserstein at $\alpha= .05$ and CPM is $r=.843$.

Based on these analyses, we conclude that using the uncorrected $\alpha= .05$ threshold is well supported, yielding high stability and high similarity with CPM.

### Virtual lesioning

In [5], the authors identify ten canonical networks of nodes: cerebellar, default mode, frontal-parietal, medial-frontal, motor, salience, subcortical, visual-association, visual I, and visual II. We tested how much each of these networks contributed to the performance of CRAM-Wasserstein and CPM by conducting virtual lesioning. That is, we altered the FC matrices of all participants to remove data about a group of canonical networks, then reran both methods on the reduced data. For each of the ten canonical networks, we performed both a held-in and a held-out virtual lesioning test. Each network contains a set of nodes, and these nodes correspond to a set of rows and columns in the FC matrix. For the held-in analysis, we *retain* only rows and columns of the FC matrix that belong to the network of interest; for the leave-out analysis, we *remove* only the nodes that belong to the network. All tests were performed using the working memory brain state in the HCP data set while predicting fluid intelligence.

Results from this analysis are in Table XXX. For the held-in lesioning, the predictive accuracy for CRAM-Wasserstein was substantially lower than with the full FC matrices. Results ranged from $r = .07$ for retaining only the visual I network to $r=.23$ for the motor network. In the held-out lesioning, results were comparable to predictive accuracy under the full FC matrices. Values ranged from $r = .38$ for the frontal-parietal network to $r=.41$. As a comparison, CRAM-Wasserstein using the full FC matrices yielded a predictive accuracy of $r = .41$. Results indicate that none of the canonical networks in isolation is responsible for the accuracy achieved when using the full matrices. Removing any single network only degraded the signal by at most $r = .03$. While using the motor network alone resulted in a predictive accuracy of $r = .23$, this result may be due to the size of the network: out of 268 nodes, 50 of them belong to the motor network, and so there may simply be more information available due to including more nodes in the analysis. Additionally, holding the motor network out only dropped the predictive accuracy to $r = .40$, indicating that whatever information is available to CRAM-Wasserstein from the motor network connections is likely present in other areas of the FC matrix.

Like CRAM-Wasserstein, CPM in the held-in analysis performed notably worse than CPM on the full data set for each canonical network, where results ranged from $r = .01$ for the cerebellar network to $r = .24$ for the salience network. In contrast to the results from CRAM-Wasserstein, the motor network did not perform particularly well, with $r = .18$. None of the held-out networks degraded the signal much, as correlations ranged from $r = .33$ for the salience network to $r = .35$ for the cerebellar network. These findings are broadly consistent with the results from the CRAM-Wasserstein lesioning analyses; no single network contains sufficient information to noticeably degrade the performance of the method when held out, but some networks alone can yield predictive correlations that are markedly better than chance.

These tests are not intended to give a full analysis of how networks affect predictive performance in the HCP working memory task; rather this section is meant to be a demonstration of how such analyses might be performed. Full lesioning analyses are beyond the scope of this paper and are a direction for future research using CRAM.

### Other metrics not considered

There are an infinite number of metrics that could be considered for purposes of the tests in this paper, and so we must necessarily limit our set of metrics. Here we describe and justify some notable exceptions from this paper.

The 𝑝-norm when 𝑝→∞ simplifies to the maximum absolute value of the elements of a vector:

$$\left\| x \right\|_{\infty}=\max_{i} \left| x_{i} \right|$$

. This norm is known as the Chebyshev or max norm. The Chebyshev distance induced by this norm was originally used in our comparisons of ID rate and predictive modeling. However, its performance was much worse than the other metrics and CPM, often being no better than chance. So, we omit the results from tests using the Chebyshev distance for simplicity.

Other distance measures induced by 𝑝-norms might have been compared, but we chose not to test them. Our results show that the Manhattan ($p=1$) and Euclidean ($p=2$) distances give approximately the same result on the ID rate and predictive modeling tests, with the Manhattan distance having the slightly better set of outcomes. Thus, we do not expect significant variation in quality for other small values of 𝑝. We also do not expect larger values of 𝑝 to improve results since the Chebyshev distance ($p\to\infty$) showed markedly worse outcomes. So, we chose to exclude other 𝑝-norm induced metrics from comparison.

Additionally, there are metrics on the space of matrices and on the space of normal distributions that were excluded due to problems with numerical stability resultant from a low ratio between the number of data points and the number of data dimensions. (Here, the number of points corresponds to the length of the timeseries used to calculate covariance between nodes—and is thus equivalent to number of TRs for a given task or rest acquisition—whereas the number of dimensions corresponds to the number of columns or rows of the connectivity matrix —and is thus equivalent to the number of nodes in a given parcellation scheme or brain atlas.) Consider the case when the number of TRs included in an fMRI scan, after motion scrubbing, is lower than the size of the atlas that has been used to parcellate the brain. In such cases the rows of the FC matrix are linearly dependent, causing the matrix to be singular (i.e., the inverse is not defined). Even when the size of the set of retained TRs is larger than the size of the atlas, if the difference between these sizes is small, then the matrix may be poorly conditioned (i.e., computing the inverse is numerically unstable). In either case, operations requiring a non-singular matrix should be avoided. Such operations include taking the inverse and dividing by the determinant. Thus, we decided to forgo tests of metrics that require FC matrices to be non-singular and well-conditioned. The set of metrics we excluded for this reason include the Bhattacharyya distance, the Hellinger distance, the Fisher-Rao metric, and the *f*-divergences. This last category includes the popular Kullback-Liebler divergence and its symmetric counterparts, the Jensen-Shannon divergence and the Jeffrey’s divergence. Further studies that include less granular atlas parcellations or larger numbers of TRs per scan would allow for the inclusion and testing of these metrics.

|  | hcp gam | hcp rest1 | hcp rest2 | hcp lang | hcp motor | hcp rela | hcp social | hcp wm | hcp emo | mean |
| --- | --- | --- | --- | --- | --- | --- | --- | --- | --- | --- |
| CRAM-Angular | 0.349 | 0.235 | 0.199 | 0.261 | 0.333 | 0.341 | 0.284 | 0.351 | 0.261 | 0.291 |
| CRAM-Euclidean | 0.348 | 0.232 | 0.171 | 0.271 | 0.325 | 0.311 | 0.301 | 0.359 | 0.270 | 0.287 |
| CRAM-Manhattan | 0.359 | 0.214 | 0.186 | 0.279 | **0.339** | 0.290 | 0.297 | 0.372 | 0.264 | 0.289 |
| CRAM-Wasserstein | **0.373** | **0.272** | **0.264** | **0.350** | 0.336 | **0.379** | **0.321** | **0.410** | **0.272** | **0.331** |
| CPM | 0.334 | 0.139 | 0.106 | 0.293 | 0.293 | 0.257 | 0.271 | 0.346 | 0.272 | 0.257 |
| KRR | 0.306 | 0.201 | 0.188 | 0.219 | 0.273 | 0.309 | 0.245 | 0.306 | 0.202 | 0.250 |

Table S1. Predictive accuracy by method and metric, as measured by Pearson’s $r$. Values are measured using 10-fold cross-validation averaged over 100 runs. Bold values indicate the most accurate result within a column. The chosen kernel for KRR is correlation between vectors, and hyperparameters are tuned using 10-fold nested cross-validation.

|  | hcp gam | hcp rest1 | hcp rest2 | hcp lang | hcp motor | hcp rela | hcp social | hcp wm | hcp emo | mean |
| --- | --- | --- | --- | --- | --- | --- | --- | --- | --- | --- |
| CRAM-Angular | 0.361 | 0.203 | 0.205 | 0.243 | 0.314 | 0.332 | 0.295 | 0.352 | 0.249 | 0.284 |
| CRAM-Euclidean | 0.355 | 0.206 | 0.183 | 0.261 | 0.308 | 0.337 | 0.315 | 0.360 | 0.254 | 0.287 |
| CRAM-Manhattan | 0.366 | 0.196 | 0.205 | 0.264 | **0.325** | 0.299 | 0.311 | 0.375 | 0.249 | 0.288 |
| CRAM-Wasserstein | **0.374** | **0.248** | **0.282** | **0.331** | 0.317 | **0.368** | **0.335** | **0.416** | 0.264 | **0.326** |
| CPM | 0.341 | 0.127 | 0.120 | 0.278 | 0.280 | 0.247 | 0.268 | 0.332 | **0.265** | 0.251 |
| KRR | 0.331 | 0.183 | 0.192 | 0.240 | 0.281 | 0.317 | 0.261 | 0.322 | 0.228 | 0.262 |

Table S2. Predictive accuracy by method and metric, as measured by Spearman’s ρ. Values are measured using 10-fold cross-validation averaged over 100 runs. Bold values indicate the most accurate result within a column. The chosen kernel for KRR is correlation between vectors, and hyperparameters are tuned using 10-fold nested cross-validation.

|  | hcp gam | hcp rest1 | hcp rest2 | hcp lang | hcp motor | hcp rela | hcp social | hcp wm | hcp emo | mean |
| --- | --- | --- | --- | --- | --- | --- | --- | --- | --- | --- |
| CRAM-Angular | 0.122 | 0.056 | 0.040 | 0.068 | 0.111 | 0.117 | 0.081 | 0.124 | 0.069 | 0.087 |
| CRAM-Euclidean | 0.121 | 0.054 | 0.029 | 0.073 | 0.106 | 0.097 | 0.091 | 0.129 | 0.073 | 0.086 |
| CRAM-Manhattan | 0.129 | 0.046 | 0.035 | 0.078 | **0.115** | 0.085 | 0.089 | 0.138 | 0.070 | 0.087 |
| CRAM-Wasserstein | **0.140** | **0.074** | **0.070** | **0.123** | 0.113 | **0.144** | **0.103** | **0.168** | **0.074** | **0.112** |
| CPM | 0.112 | 0.020 | 0.012 | 0.080 | 0.086 | 0.041 | 0.074 | 0.112 | 0.074 | 0.068 |
| KRR | 0.093 | 0.040 | 0.035 | 0.048 | 0.074 | 0.095 | 0.060 | 0.093 | 0.041 | 0.065 |

Table S3. Predictive accuracy by method and metric, as measured by coefficient of determination. Values are measured using 10-fold cross-validation averaged over 100 runs. Bold values indicate the most accurate result within a column. The chosen kernel for KRR is correlation between vectors, and hyperparameters are tuned using 10-fold nested cross-validation.

|  | hcp gam | hcp rest1 | hcp rest2 | hcp lang | hcp motor | hcp rela | hcp social | hcp wm | hcp emo | mean |
| --- | --- | --- | --- | --- | --- | --- | --- | --- | --- | --- |
| CRAM-Angular | 4.20 | 4.38 | 4.43 | 4.38 | 4.21 | 4.21 | 4.32 | 4.20 | 4.34 | 4.29 |
| CRAM-Euclidean | 4.20 | 4.37 | 4.46 | 4.34 | 4.22 | 4.26 | 4.27 | 4.17 | 4.31 | 4.29 |
| CRAM-Manhattan | 4.17 | 4.39 | 4.43 | 4.32 | **4.19** | 4.30 | 4.27 | 4.14 | 4.32 | 4.28 |
| CRAM-Wasserstein | **4.13** | **4.28** | **4.30** | **4.16** | **4.19** | **4.12** | **4.21** | **4.06** | **4.28** | **4.19** |
| CPM | 4.27 | 4.63 | 4.74 | 4.33 | 4.35 | 4.43 | 4.40 | 4.24 | 4.41 | 4.42 |
| KRR | 4.24 | 4.42 | 4.47 | 4.35 | 4.28 | 4.32 | 4.32 | 4.24 | 4.36 | 4.33 |

Table S4. Predictive accuracy by method and metric, as measured by root mean squared error. Values are measured using 10-fold cross-validation averaged over 100 runs. Bold values indicate the most accurate result within a column. The chosen kernel for KRR is correlation between vectors, and hyperparameters are tuned using 10-fold nested cross-validation.

|  | hcp gam | hcp rest1 | hcp rest2 | hcp lang | hcp motor | hcp rela | hcp social | hcp wm | hcp emo | mean |
| --- | --- | --- | --- | --- | --- | --- | --- | --- | --- | --- |
| CRAM-Angular | 0.089 | 0.042 | 0.078 | 0.252 | 0.190 | 0.127 | 0.104 | 0.169 | 0.200 | 0.139 |
| CRAM-Wasserstein | **0.149** | 0.077 | **0.081** | **0.362** | **0.212** | **0.218** | **0.219** | **0.233** | **0.257** | **0.201** |
| CPM | 0.077 | **0.078** | 0.000 | 0.266 | 0.157 | 0.122 | 0.047 | 0.089 | 0.194 | 0.114 |

Table S5. Clustering coefficients of identified networks by method and metric. Values are measured using 10-fold cross-validation averaged over 100 runs. Bold values indicate the largest result within a column.

| Dim | hcp gam | hcp rest1 | hcp rest2 | hcp lang | hcp motor | hcp rela | hcp social | hcp wm | hcp emo | mean |
| --- | --- | --- | --- | --- | --- | --- | --- | --- | --- | --- |
| 50% | 0.365 | 0.222 | 0.196 | 0.318 | 0.327 | 0.335 | 0.329 | 0.378 | 0.273 | 0.305 |
| 60% | 0.363 | 0.258 | 0.238 | 0.316 | 0.333 | 0.344 | 0.321 | 0.378 | 0.273 | 0.314 |
| 70% | 0.371 | 0.263 | 0.261 | 0.328 | 0.337 | 0.346 | 0.320 | 0.392 | 0.270 | 0.321 |
| 80% | 0.372 | 0.267 | 0.268 | 0.329 | 0.339 | 0.348 | 0.324 | 0.399 | 0.272 | 0.324 |
| 90% | 0.373 | 0.268 | 0.267 | 0.341 | 0.340 | 0.366 | 0.322 | 0.405 | 0.273 | 0.328 |
| 100% | 0.373 | 0.272 | 0.264 | 0.350 | 0.336 | 0.379 | 0.321 | 0.410 | 0.272 | 0.331 |
| 110% | 0.380 | 0.275 | 0.259 | 0.351 | 0.334 | 0.389 | 0.326 | 0.413 | 0.271 | 0.333 |
| 120% | 0.388 | 0.274 | 0.262 | 0.351 | 0.331 | 0.395 | 0.324 | 0.414 | 0.268 | 0.334 |
| 130% | 0.387 | 0.272 | 0.263 | 0.348 | 0.328 | 0.400 | 0.328 | 0.412 | 0.268 | 0.334 |
| 140% | 0.387 | 0.267 | 0.270 | 0.347 | 0.327 | 0.399 | 0.327 | 0.413 | 0.264 | 0.333 |
| 150% | 0.388 | 0.261 | 0.280 | 0.344 | 0.326 | 0.395 | 0.327 | 0.409 | 0.259 | 0.332 |

Table S6. Predictive correlation of HCP fluid intelligence score from the HCP working memory task using CRAM-Wasserstein vs. MDS dimension

Figure S1

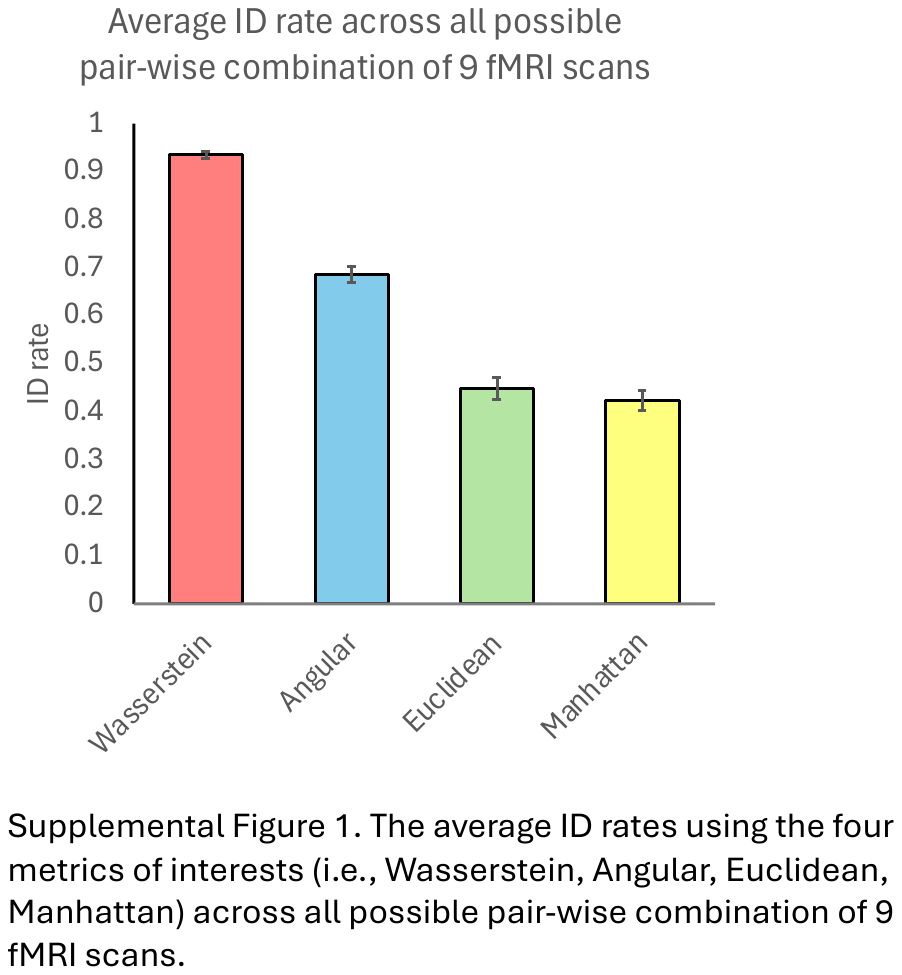

Figure S2

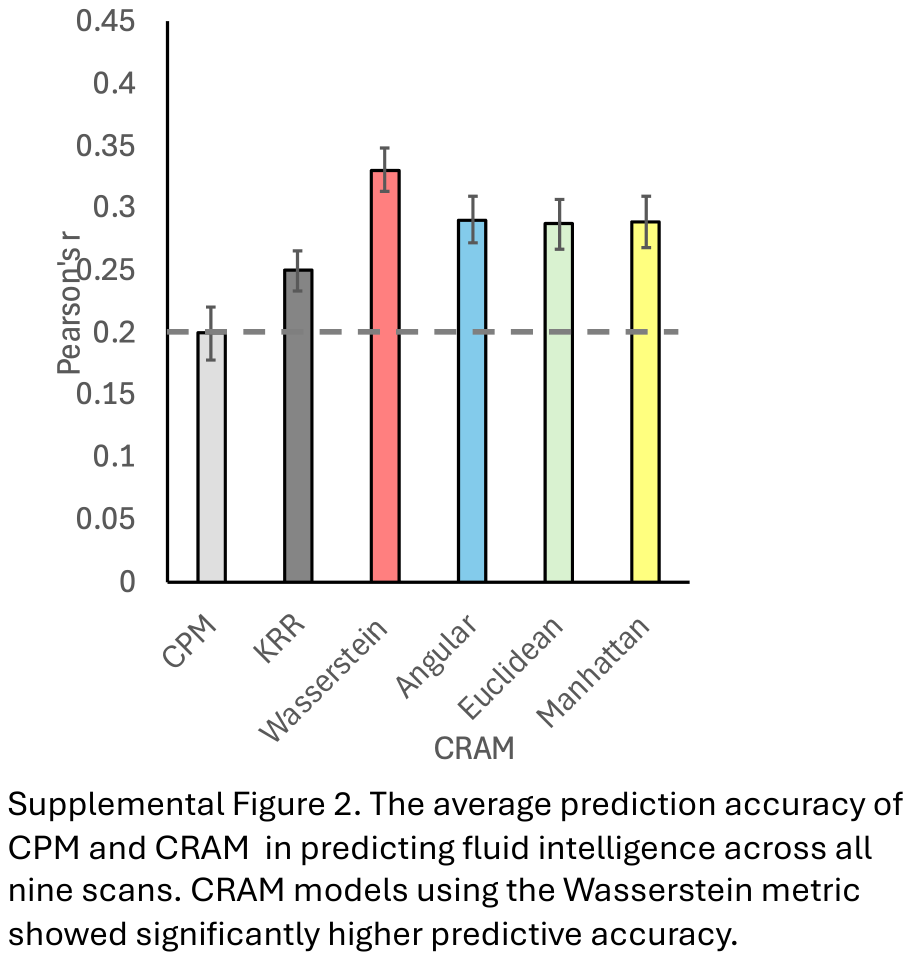

Figure S3

Predictive correlation vs. embedding dimension

Average r

| Number dimensions as a fraction rule-of-thumb | average r |
| --- | --- |
| 50% | 0.304611 |
| 60% | 0.313849 |
| 70% | 0.321151 |
| 80% | 0.324189 |
| 90% | 0.328172 |
| 100% | 0.330887 |
| 110% | 0.333131 |
| 120% | 0.334133 |
| 130% | 0.33384 |
| 140% | 0.333309 |
| 150% | 0.332201 |

Supplemental Figure 3. Predictive correlation using CRAM-Wasserstein vs. MDS dimension
